# Metformin attenuates hyperglycaemia-stimulated pro-fibrotic gene expression in vascular adventitial fibroblasts via inhibition of Discoidin Domain Receptor 2

**DOI:** 10.1101/2022.10.24.513528

**Authors:** Allen Sam Titus, Mereena George Ushakumary, Harikrishnan Venugopal, Mingyi Wang, Edward G. Lakatta, Shivakumar Kailasam

## Abstract

Molecular mechanisms underlying the diverse therapeutic effects of anti-diabetic metformin, beyond its anti-hyperglycaemic effects, remain largely unclear. Metformin is reported to reduce the long-term complications of diabetes, including cardiovascular fibrosis and remodelling. Our recent investigations show that Discoidin Domain Receptor 2 (DDR2), a collagen receptor tyrosine kinase, has an obligate regulatory role in collagen type I gene expression in cardiac and vascular adventitial fibroblasts, and that it may be a molecular link between arterial fibrosis and metabolic syndrome in rhesus monkeys. Using gene knockdown and over-expression approaches, the present study examined whether DDR2 is a target of metformin, and whether, by targeting DDR2, it inhibits fibronectin and collagen expression in vascular adventitial fibroblasts exposed to hyperglycaemic conditions. Metformin was found to attenuate hyperglycaemia-induced increase in DDR2 mRNA and protein expression by inhibiting TGF-β1/ Smad2/3 signalling that mediates the stimulatory effect of hyperglycaemia on DDR2 expression. Metformin also inhibited DDR2-dependent expression of fibronectin and collagen, indicating that it regulates these matrix proteins via DDR2 inhibition. The findings identify DDR2, a major mediator of cardiovascular remodelling, as a molecular target of metformin, thereby uncovering the molecular basis of its protective role in vascular fibrosis, and possibly, cardiac fibrosis associated with diabetic cardiomyopathy.

## Introduction

Metformin is a widely used anti-diabetic drug whose beneficial effects far exceed its anti-hyperglycaemic action and are possibly mediated by mechanisms distinct from its metabolic actions such as inhibition of mitochondrial enzymes, and consequent activation of AMP-activated protein kinase [1,2]. Several studies point to its protective role in cancer, polycystic ovary syndrome, and neurological disorders [3–6]. Metformin is reported to exert potent anti-fibrotic effects in the lung by modulating metabolic pathways, inhibiting TGF-β1 action, suppressing collagen production, activating PPARγ signalling and inducing lipogenic differentiation in myofibroblasts [7]. A number of clinical studies have shown that metformin reduces the incidence of cardiovascular events and associated mortality [8,9]. It has been demonstrated that metformin mitigates cardiac fibrosis induced by pressure overload and inhibits collagen synthesis in cardiac fibroblasts via inhibition of the TGF-β1/Smad3 signalling pathway [10]. Another study showed that the drug inhibits inflammatory responses through suppression of the NF-κB inflammatory signalling pathway, improves vascular endothelial function, and inhibits cardiomyocyte apoptosis and myocardial fibrosis [9]. Notably, metformin protects against adverse fibrotic vascular remodelling [9], which is a major complication of diabetes that contributes to end-organ damage. Notwithstanding these reports, the precise mechanisms underlying the diverse therapeutic effects of metformin remain largely elusive [2]. Admittedly, it is of immense scientific interest and clinical relevance to identify the molecular targets of metformin and delineate the mechanisms that underlie its widely acknowledged cardiovascular protective effects, especially its anti-fibrotic action.

In this regard, several studies have shown that Discoidin Domain Receptor 2 (DDR2), a collagen receptor tyrosine kinase, is a major regulator of a wide array of cellular processes such as extracellular matrix (ECM) turnover, proliferation, differentiation, migration and survival in different cell types, including cardiac fibroblasts and vascular cells [11–17]. Further, our recent studies provide robust evidence that DDR2 may be a molecular link between arterial fibrosis and metabolic syndrome in rhesus monkeys [18]. The predominant localisation of DDR2 in cells of mesenchymal origin and its indispensable role in collagen production in cardiovascular cells make it a potential player, and hence a drug target, in adverse cardiovascular tissue fibrosis and remodelling associated with pathological states such as diabetes and atherosclerosis.

The present study examined whether DDR2 is a target of metformin, and whether, by targeting DDR2, the drug inhibits fibronectin and collagen expression in vascular adventitial fibroblasts exposed to hyperglycaemic conditions. We demonstrate, for the first time, that metformin prevents hyperglycaemia-induced increase in DDR2 expression in vascular adventitial fibroblasts by inhibiting the TGF-β1/ Smad2/3 signalling pathway, which enhances DDR2 expression in response to hyperglycaemia. Additionally, we also show that inhibition of DDR2 by metformin results in inhibition of DDR2-dependent fibronectin and collagen type I expression in these cells. The findings are particularly important insofar as they identify DDR2, a major mediator of cardiovascular remodelling [11,12,18], as a molecular target of metformin, uncovering the molecular basis of its protective role in vascular fibrosis, and possibly, cardiac fibrosis associated with diabetic cardiomyopathy.

## Results and Discussion

The incidence of diabetes mellitus is on the rise globally due to a variety of reasons, including increasing prevalence of obesity, genetic susceptibility, and ageing [19]. Since conventional treatment modalities aiming at rigorous glycaemic control may not completely eliminate cardiovascular risk burden, other strategies targeting critical pathogenetic mechanisms may offer hope. In this regard, it is pertinent to note that one of the hallmarks of vascular complications associated with diabetes is abnormal matrix deposition by vascular adventitial fibroblasts and vascular smooth muscle cells, leading to vascular fibrosis and remodelling of the ECM within the vascular wall [20]. This is integral to intimal-medial thickening, reduced lumen diameter and compromised vascular resilience [9]. Hyperglycaemia-induced alterations in vascular wall structure may contribute to the pathogenesis of microvascular and macrovascular complications attributed to diabetes, resulting eventually in impairment of target organs such as the heart, kidneys and brain. Targeting mechanisms that regulate ECM gene expression in vascular cells is clearly a promising option to prevent the cardiovascular complications of diabetes. This study focused on DDR2 as a target of metformin, examined the mechanism by which metformin inhibits DDR2, and in light of our earlier published studies, explored the consequences of DDR2 inhibition by metformin vis-a-vis matrix protein expression.

Exposure of adventitial fibroblasts to high glucose at 30 mM (HG) was found to cause a significant increase in fibronectin (Figure 1) and collagen protein expression (Figure 2). HG also caused a significant increase in DDR2 mRNA and protein levels (Figures 3 A and B). Our previous studies with mannitol at the same concentration had ruled out the possibility that these effects of HG may be osmolarity-related [18]. To ascertain whether DDR2 may have a role in HG-stimulated fibronectin expression in adventitial fibroblasts, the cells were transfected with DDR2 siRNA, as described under Methods, and after confirming knockdown, fibronectin expression was determined following exposure of the cells to HG. DDR2 knockdown attenuated the stimulatory effect of HG on fibronectin expression (Figure 4). In tandem with earlier observations [11–14,21], the finding shows conclusively that DDR2 has an indispensable regulatory role in the expression of fibronectin and collagen type 1 in both cardiac fibroblasts and vascular adventitial fibroblasts. The shared characteristic of a regulatory relationship between DDR2 and the two major ECM proteins in cardiac and vascular cells that express these proteins may be an important determinant of fibrosis in the cardiovascular system. Admittedly, any drug that targets DDR2 would impact the expression of these key matrix proteins in cardiovascular cells and mitigate the cardiovascular complications of diabetes, which affect the prognosis of patients.

**Figure 1:**
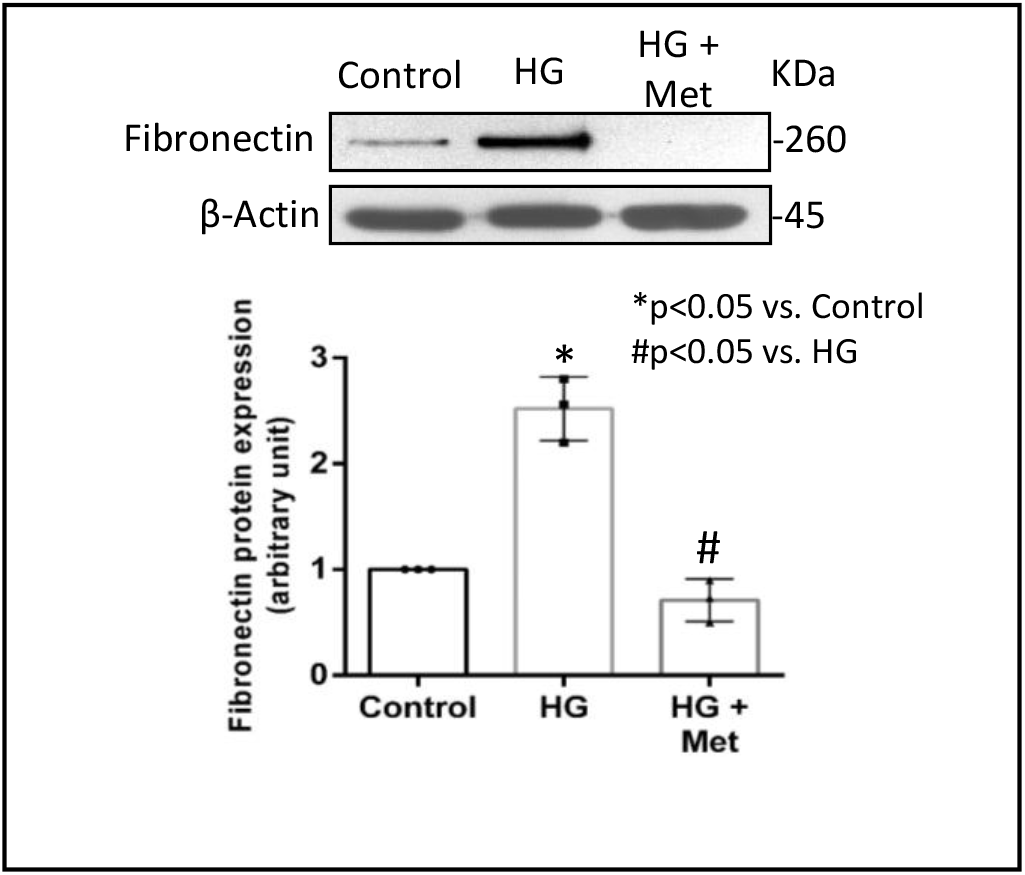
Metformin prevents HG-induced increase in fibronectin protein expression in vascular adventitial fibroblasts. Cells were pre-treated with metformin hydrochloride (10 μM) for 45 mins prior to HG supplementation. Protein was isolated after 12 h of HG and subjected to western blot analysis for detection of fibronectin, with β-actin as loading control. *p< 0.05 vs. control, #p< 0.05 vs. HG. Data are representative of 3 independent experiments, n=3. Error bars represent Standard Error (SE).

**Figure 2:**
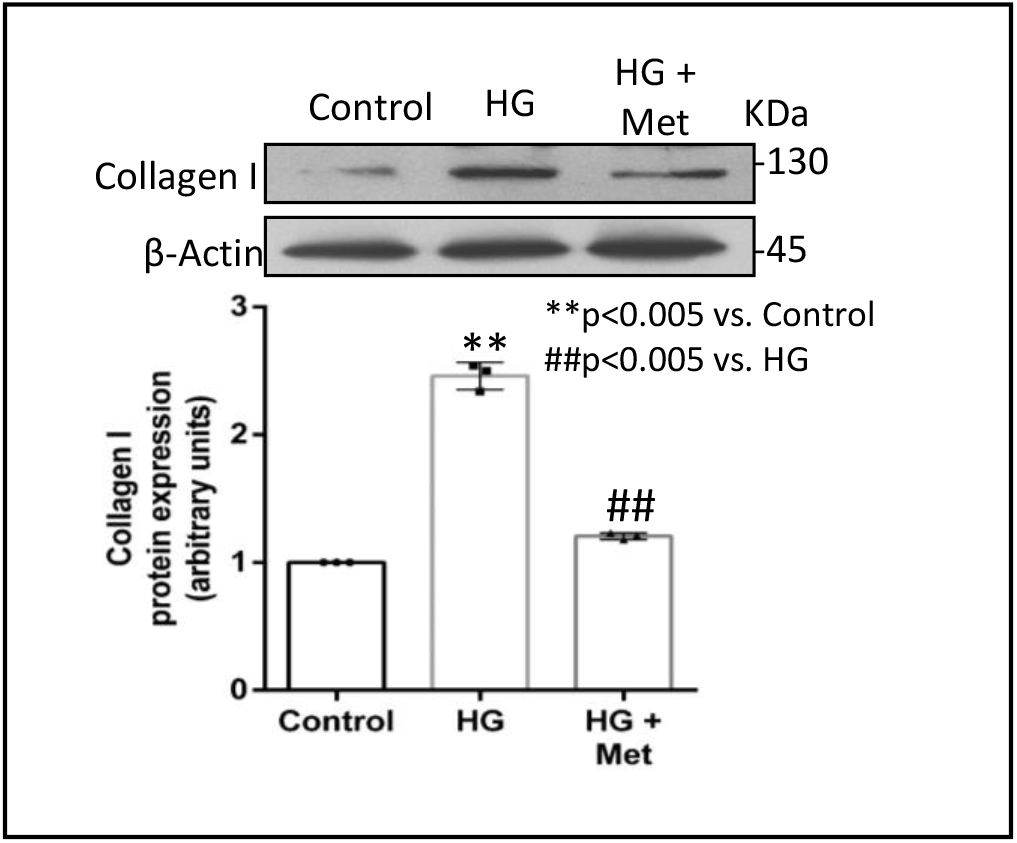
Metformin prevents HG-induced increase in collagen type I protein expression in vascular adventitial fibroblasts. Cells were pre-treated with metformin hydrochloride (10 μM) for 45 mins prior to HG supplementation. Protein was isolated after 12 h of HG and subjected to western blot analysis for detection of collagen type I, with β-actin as loading control. **p< 0.005 vs. control, ##p< 0.005 vs. HG. Data are representative of 3 independent experiments, n=3. Error bars represent Standard Error (SE).

**Figure 3:**
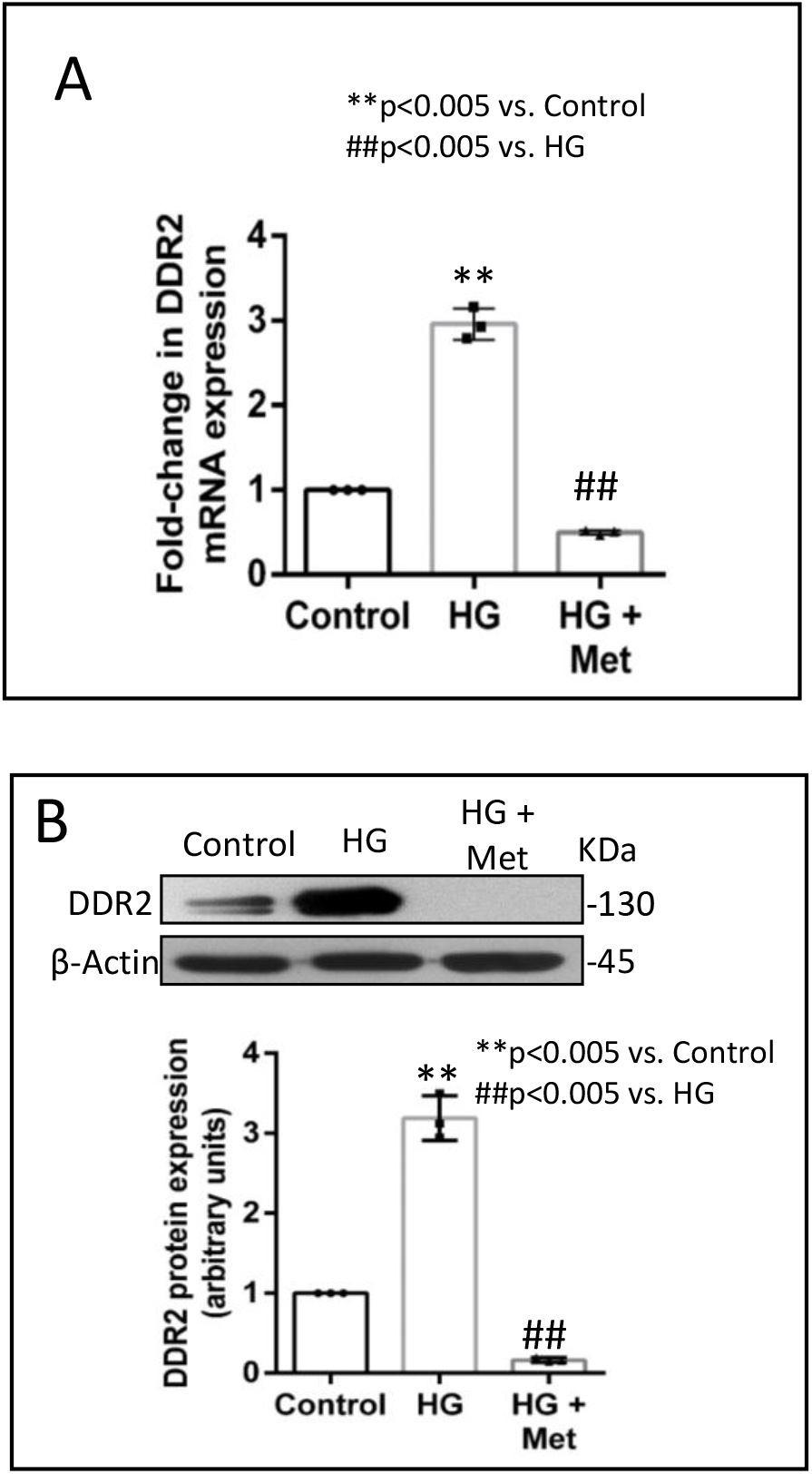
Metformin prevents HG-induced increase in DDR2 expression in vascular adventitial fibroblasts. Cells were pre-treated with metformin hydrochloride (10 μM), 45 mins prior to HG supplementation. **(A)** DDR2 mRNA levels were determined by Taqman real-time PCR at 6 h post-HG treatment, with β-actin as loading control. **p< 0.005 vs. control, ##p< 0.005 vs. HG (One-way ANOVA p< 0.05). **(B)** Protein was isolated after 12h of HG treatment and subjected to western blot analysis for detection of DDR2, with β-actin as loading control. **p< 0.005 vs. control, ##p< 0.005 vs. HG. Data are representative of 3 independent experiments, n=3. Error bars represent Standard Error (SE).

**Figure 4:**
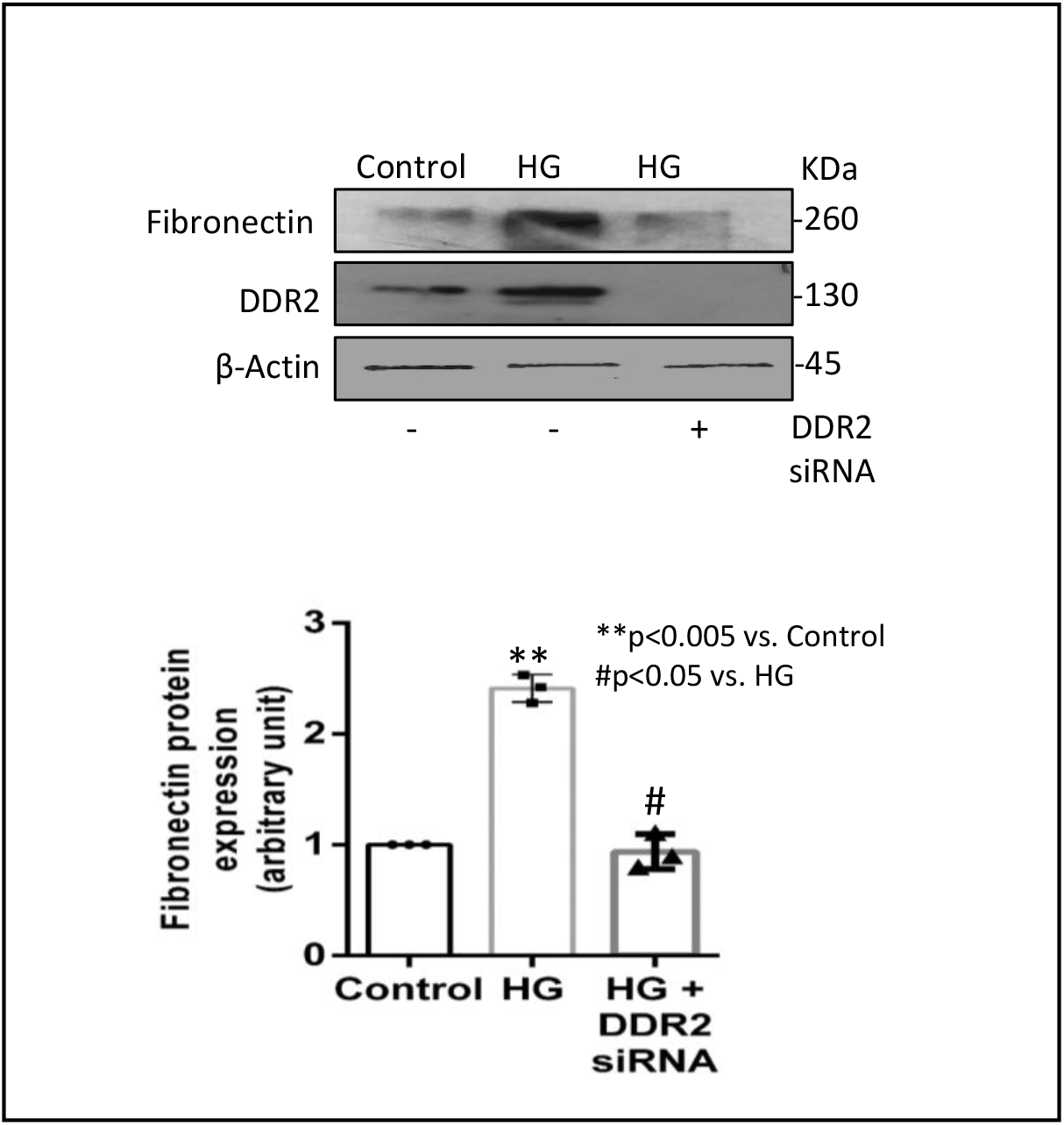
DDR2 knockdown prevents HG-induced increase in fibronectin expression in vascular adventitial fibroblasts. Vascular adventitial fibroblasts were transiently transfected with DDR2 siRNA or scrambled siRNA. Following exposure of the transfected cells to HG for 12 h, fibronectin protein expression was examined by western blot analysis and normalized to β-actin. **p< 0.005 vs. control, #p< 0.05 vs. HG. Data are representative of 3 independent experiments, n=3. Error bars represent Standard Error (SE).

Concurrent treatment of vascular adventitial fibroblasts with metformin was found to significantly reduce HG-induced increase in fibronectin and collagen expression in these cells (Figures 1 and 2). Notably, metformin also reduced DDR2 expression in HG-treated cells (Figures 3 A and B), indicating that it may regulate fibronectin and collagen via a DDR2-dependent mechanism in vascular adventitial fibroblasts. Both collagen type I and fibronectin are importantly involved in the regulation of cardiovascular pathophysiology. Although earlier studies focused predominantly on collagen, fibronectin is now implicated in the pathogenesis of pulmonary and vascular fibrosis. Inhibition of fibronectin as a strategy to diminish fibrosis and improve cardiac function has been demonstrated in an animal model of heart failure [22]. Polymerized fibronectin influences the deposition, maturation and stability of other ECM proteins, including collagen type I, which makes it an important regulator of ECM [22]. Therefore, targeted regulation of collagen and fibronectin expression in cardiovascular cells, or a common regulator of these two important ECM proteins, would provide a novel therapeutic approach to prevent or reduce cardiovascular fibrosis. The inhibitory action of metformin on DDR2 reported here needs to be viewed in light of these observations.

Next, we examined how metformin may inhibit DDR2 expression, focusing on the role of Transforming Growth Factor-β 1 (TGF-β 1) that is reported to be involved in the progression of many diseases, including cardiovascular diseases in which it is associated with fibrosis, epithelial-to-mesenchymal transition and inflammation [23,24]. In the present study, hyperglycaemia was found to enhance TGF-β1 mRNA and protein expression in vascular adventitial fibroblasts (Figures 5 A and B). Inhibition of TGF-β1 using SB 431542 or TGF-β1 siRNA, and Smad 2/3 inhibition using PD169316 or SIS3, abolished HG-induced increase in fibronectin expression in these cells (Figures 6 A and B). Interestingly, while TGF-β1 siRNA abolished HG-induced increase in DDR2 expression (Figure 6 C), consistent with the previously reported regulatory role of TGF-β1 in DDR2 expression [18], over-expression of DDR2 in TGF-β1-silenced cells restored fibronectin, which showed that DDR2 acts downstream of TGF-β1 to regulate fibronectin expression (Figure 6 C). Notably, metformin was found to abolish HG-stimulated increase in TGF-β1 mRNA and protein expression (Figures 5 A and B), indicating that suppression of TGF-β1/Smad 2/3 signalling underlies the negative regulatory effect of the drug on DDR2.

**Figure 5:**
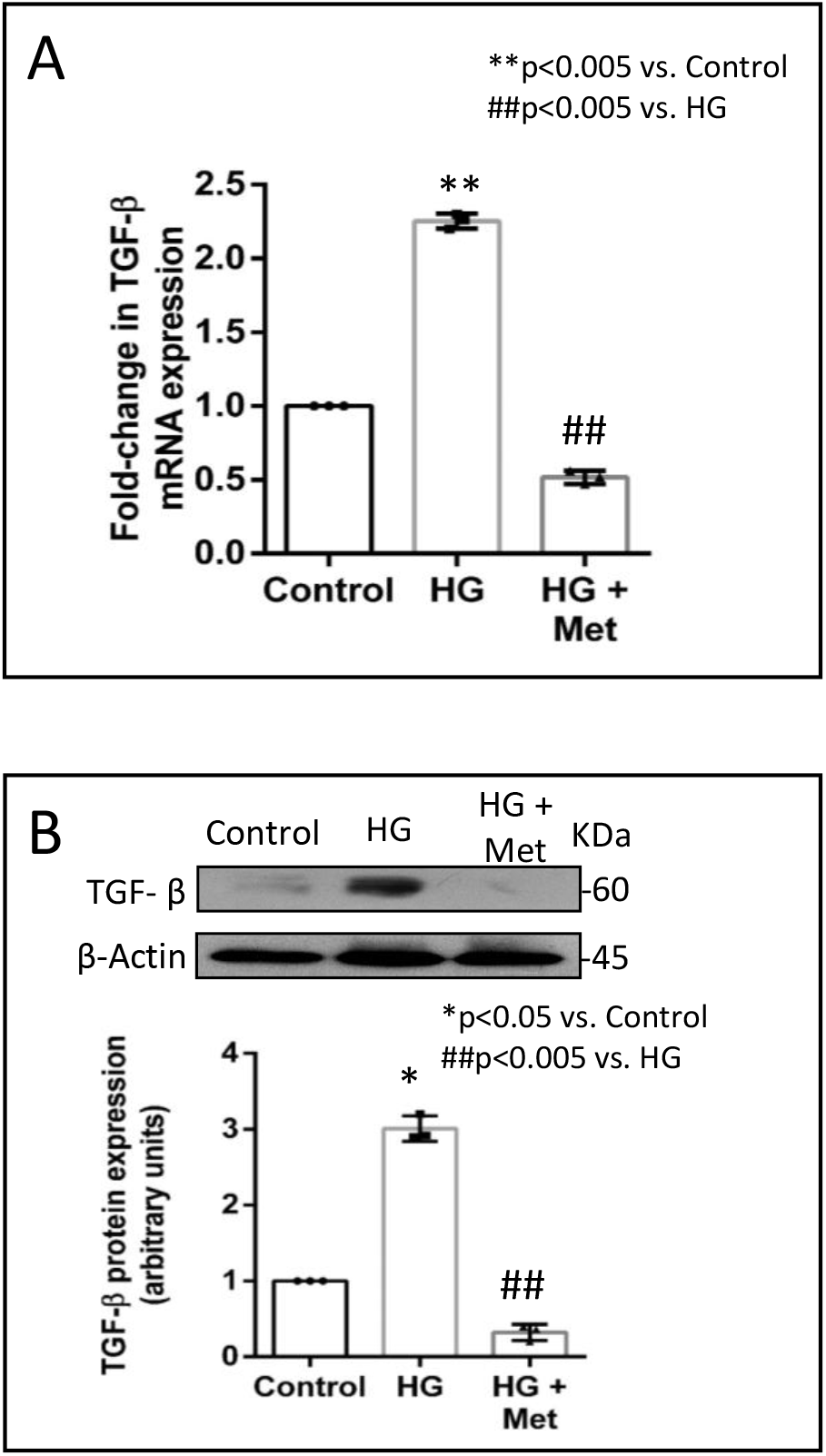
Metformin prevents HG-induced increase in TGF-β1 mRNA (A) and protein (B) expression in vascular adventitial fibroblasts. Cells were pre-treated with metformin hydrochloride (10 μM), 45 mins prior to HG supplementation. **(A)** TGF-β1 mRNA levels were determined by Taqman real-time PCR at 6 h post-HG treatment, with β-actin as loading control. **p< 0.005 vs. control, ##p< 0.005 vs. HG. **(B)** Protein was isolated after 12 h of HG and subjected to western blot analysis for detection of TGF-β1, with β-actin as loading control. *p< 0.05 vs. control, ##p< 0.005 vs. HG. Data are representative of 3 independent experiments, n=3. Error bars represent Standard Error (SE).

**Figure 6:**
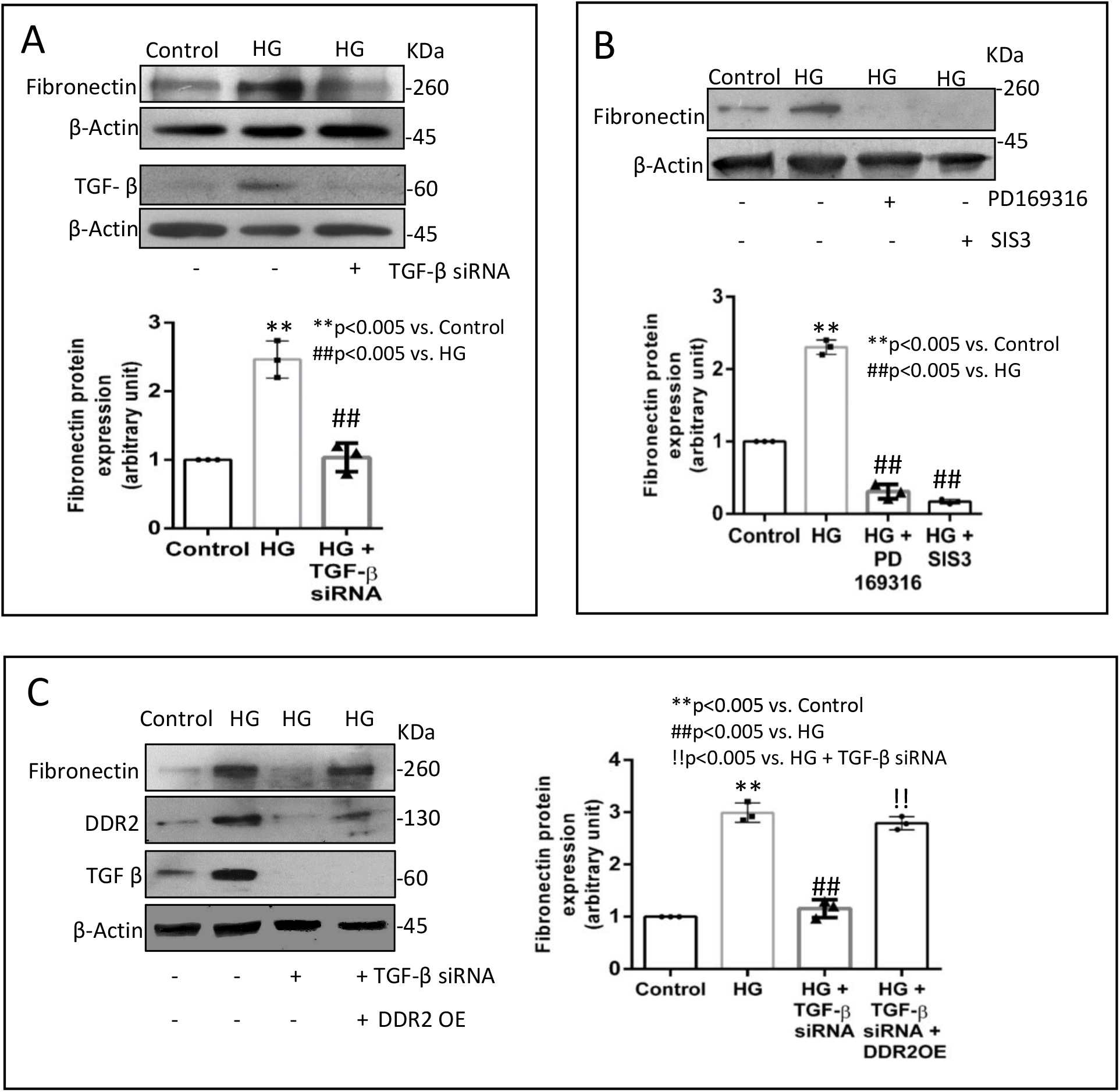
DDR2 acts downstream of TGF-β1 to regulate fibronectin expression. **(A)** Vascular adventitial fibroblasts were transiently transfected with TGF-β1 siRNA 1 and 2 (5 pmol each) or scrambled siRNA prior to treatment with HG for 12 h. Protein was isolated and subjected to western blot analysis for detection of fibronectin, with β-actin as loading control. Validation of TGF-β1 knockdown is also shown. **p< 0.005 vs. control, ##p< 0.005 vs. HG and ns -not significant vs. control. **(B)** Sub-confluent, quiescent cultures of vascular adventitial fibroblasts in M199 were pre-treated with SMAD2 inhibitor, PD169316, or SMAD3 inhibitor, SIS3, for 1 h and subsequently with HG for 12 h. Following treatment with HG, protein was isolated and subjected to western blot analysis for detection of fibronectin, with β-actin as loading control. **p< 0.005 vs. control, ##p< 0.005 vs. HG and ns - not significant vs. control. **(C)** Vascular adventitial fibroblasts were co-transfected with DDR2 plasmid (1μg) and TGF-β1 siRNA 1 and 2 (5 pmol each) (lipofectamine 2000-complexed) for 8 h in OptiMEM. After a recovery period of 12 h, the cells were subjected to HG for 12 h. Following treatment with HG, protein was isolated and subjected to western blot analysis for detection of fibronectin, with β-actin as loading control. Validation of DDR2 OE and TGF-β1 knockdown is also shown. **p<0.005 vs. Control, ##p<0.005 vs. HG, !!p<0.005 vs. HG + TGF-β siRNA. Data are representative of 3 independent experiments, n=3. Error bars represent Standard Error (SE).

While discussing the action of metformin in the context of hyperglycaemia, it is pertinent to appreciate the reported link between hyperglycaemia and reactive oxygen species (ROS) whose role in mediating both physiological and pathophysiological signal transduction profoundly impacts gene expression in diverse cellular contexts. Many studies report that high glucose concentrations promote increased ROS generation, leading to oxidative stress and consequent cellular changes that underlie the complications of diabetes, including cardiovascular diseases [25]. High glucose concentrations are also reported to enhance intracellular synthesis of Angiotensin II (Ang II) in cardiac myocytes, vascular smooth muscle cells and cardiac fibroblasts [26]. Moreover, in cardiac fibroblasts, Ang II acts via ROS to enhance DDR2 expression via the redox-sensitive transcription factor, NF-κB [11]. Remarkably, several studies report the inhibitory effect of metformin on ROS and oxidative stress [27]. It is therefore possible that metformin may negatively impact ROS signalling to inhibit DDR2 and DDR2-dependent downstream events.

In conclusion, this study expands our understanding of the mode of action of metformin, throwing light on a novel mechanism by which this drug of choice in the treatment of diabetes may prevent excessive matrix accumulation in the vessel wall and protect against vascular fibrosis and dysfunction. The inhibitory effect of metformin on TGF-β1-dependent DDR2 expression and consequent down-regulation of DDR2 signalling is an important finding with immense clinical implications because of the characteristic presence of DDR2 in cardiac fibroblasts and vascular cells and its critical role in the regulation of these cells. The obligate role of DDR2 in collagen and fibronectin expression in cardiac and vascular adventitial fibroblasts and its association with arterial fibrosis establish its role as an important determinant of fibrosis in the cardiovascular system [11,12,18,21]. DDR2 also regulates the phenotypic transformation of Angiotensin II-stimulated cardiac fibroblasts [12], and is a positive regulator of cardiac fibroblast proliferation [28]. It confers apoptosis resistance on cardiac fibroblasts [28], which may underlie the role of these cells in adverse myocardial fibrosis following injury. Importantly, DDR2 regulates the expression of the Ang II receptor, AT1, in cardiac fibroblasts [21], which would impact the pleiotropic actions of Ang II on cardiovascular cells. In this context, it is particularly important to note that metformin is also reported to inhibit all of these processes that are under the regulatory control of DDR2. Thus, in different experimental models, metformin has been shown to inhibit the phenotypic transformation of Ang II-stimulated cardiac fibroblasts [29] and negatively regulate the expression of collagen, fibronectin [30], and the AT1 receptor [31], and inhibit cardiac fibrosis via inhibition of the TGF-β1/Smad3 signalling pathway [10,32]. What is, however, not clear from these studies is whether a single critical regulator of multiple pathways could mediate the cardiovascular protective effects of metformin. Pending in vivo and clinical studies, it seems very likely that the manifold cardiovascular benefits of metformin action are attributable, at least in part, to the suppression of DDR2 signalling.

## Materials and Methods

PD169316 and SIS3 were from Calbiochem (San Diego, USA). Collagenase type II and elastase (lyophilized) were from Worthington (Columbus, USA). Glucose, metformin hydrochloride (Cat No. 1396309), SB431542, M199 and TGF-β1 (Cat No. T7039) were obtained from Sigma-Aldrich (St. Louis, MO, USA). Lipofectamine 2000 was from Invitrogen (Carlsbad, CA, USA). The Chemiluminescence Western blot detection reagent was from Thermo Fisher Scientific (Waltham, MA, USA). DDR2, TGF-β1 siRNA and control siRNAs were from Ambion (Foster City, CA, USA). Fibronectin siRNA was custom-synthesized by Eurogentec (Liège, Belgium). The rat DDR2/CD167b Gene ORF cDNA clone expression plasmid was obtained from Sino Biologicals (Beijing, China). Opti-MEM and Fetal Bovine serum (FBS) were from GIBCO (Waltham, MA, USA). All cell culture ware was purchased from BD Falcon (Corning, NY, USA). Primary antibody against DDR2 (Cat No. 12133S) and TGF-β1 (Cat No. 3711) were obtained from Cell Signalling Technology (Danvers, MA, USA). Primary antibodies against fibronectin (Cat No. sc9068) and collagen I (Cat No. sc293182) were from Santa Cruz Biotechnology (Dallas, TX, USA). Loading control β-actin (Cat No. A2228) antibody was obtained from Sigma-Aldrich. All antibodies were used after dilution (1:1000). XBT X-ray Film was from Carestream (Rochester, NY, USA).

### Isolation of rat vascular adventitial fibroblasts

Vascular adventitial fibroblasts were isolated from young adult male Sprague-Dawley rats following protocols previously standardized by us [18]. Aorta was removed from left subclavian origin, washed in Hank’s Balanced Salt Solution (HBSS) solution, and then transferred to clean 100 mm dish with fresh HBSS in a sterile hood. Fat was removed gently under a microscope and incubated in a 35 mm dish containing 2 ml Digestion Medium I for 30 min at 37°C. Tissue was then placed in fresh HBSS, cut open, separated into the outermost adventitial layer after scraping out the endothelium, and incubated in Digestion Medium II for 90 min at 37°C. After incubation, an equal volume of Medium-199 (M199) containing 10% FBS was added to the dish and mixed well for proper dissociation. Cells were collected by filtering through a cell strainer (70 μm) and centrifuged at 1500 rpm for 5 min. The cell pellet was collected and resuspended in 0.5 ml of M199 containing 10% FBS and seeded on 12-well plate. At 24 h after isolation, the supernatant containing unattached cells was discarded and the cells were incubated with M199 containing 10% FBS in a CO_2_ incubator at 37°C. Cultures of adventitial fibroblasts were characterized by immunocytochemistry and checked for cross-contamination. Adventitial fibroblasts stained positive for DDR2 and negative for Desmin, as shown earlier [18]. Cells from passage 3–4 were used for the experiments.

### High glucose and metformin treatment

Vascular adventitial fibroblasts were grown in Dulbecco’s Modified Eagle Medium (DMEM) (5.5 mM glucose, normal medium) for 24 h prior to the experiments. D-glucose was then supplemented to achieve a final concentration of 30 mM D-glucose (high glucose (HG); hyperglycaemic media) and the cells were incubated for 6 h (for mRNA expression) and 12 h (for protein expression). Mannitol at 30 mM served as osmotic control. In the metformin group, metformin hydrochloride was added to the normal medium for a final concentration of 10 μM, 45 mins prior to high glucose supplementation.

### Western blot analysis

Sub-confluent cultures of adventitial fibroblasts were subjected to the indicated treatments for 12 h, following which relative protein expression was determined by western blot analysis by standard protocols, using β-actin as loading control. Enhanced chemiluminescence reagent was used to detect the proteins. Protein expression was quantified by densitometric analysis of scanned films using ImageJ software [33].

### Real-time polymerase chain reaction analysis

Sub-confluent cultures of adventitial fibroblasts were subjected to the indicated treatments and total RNA was isolated using TRI reagent, following the manufacturer’s instructions. After DNase I treatment, 2 μg of total RNA was reverse transcribed to cDNA with random primers and M-MLV reverse transcriptase. TaqMan quantitative RT-PCR analysis was performed using the ABI prism 7500 Sequence Detection System (Applied Biosystems,CA) with specific FAM-labeled probes. PCR reactions were carried out over 40 cycles, as per the manufacturer’s instructions. DDR2 and collagen type I mRNA expression levels were normalized to β-actin and 18s rRNA, respectively.

### RNA interference

The gene knockdown protocol by RNA interference was as reported by us earlier (18). Briefly, cells were seeded on 12-well plates at 8×10^4^ cells/well. After 24 h, the cells were incubated in Opti-MEM with Ambion pre-designed Silencer Select siRNA [5 pmol DDR2, 5 pmol TGF-β1(siRNA 1 and 2) or scrambled siRNA, control] and Lipofectamine 2000 (2 μl) for 19 h. After an additional incubation in M199 with 10% FBS for 12 h, the cells were treated with high glucose for the indicated duration. Cell lysate was prepared in SDS lysis buffer, denatured, and used for western blot analysis.

### Over-expression of DDR2

Strong constitutive expression of DDR2 was achieved, as previously described by us [18], under a pCMV promoter, purchased as cDNA clone from Sinobiologicals, China (Rat DDR2/CD167b Gene ORF cDNA). The DDR2 clone size was verified by PCR amplification of the primers in the kit. The size of the plasmid was analyzed by single site restriction digestion at the Mlu-1 site. DDR2 overexpression plasmid cocktail was prepared with 1 μg plasmid and 3 μl Lipofectamine 2000 in 100 μl Opti-MEM, and added to the cultures for incubation in M199+10% FBS for 8 hours.

### Statistical Analysis

Data are expressed as Mean ± SEM (Standard Error of Mean). Statistical analysis was performed using Student’s *t*-test (unpaired,2-tailed) for comparisons involving 2 groups. For comparisons involving more than 2groups, the data were analyzed by one-way ANOVA (with one variable). *p*≤ 0.05 was considered significant. The in vitro data presented are representative of 3 or 4 independent experiments.

## Abbreviations

DDR2: Discoidin Domain Receptor 2
TGF-β1: Transforming Growth Factor-Beta 1
Smad2/3: SMAD Family Member 2/3
PPARγ: Peroxisome proliferation-activated receptor gamma
NF-κB: Nuclear factor-kappa Beta
Ang-II: Angiotensin II
AT1: Angiotensin II receptor type 1

## Author Contributions

AST, SK, HV, MGU, EGL and MW conceived the study; AST and SK designed the study; AST, HV and MGU performed the experiments; AST and SK analyzed the data; AST and SK interpreted the results of the experiments; AST prepared the figures; AST and SK drafted the manuscript; All authors have read and agreed to the published version of the manuscript.

## Funding

SK was supported by the Indian Council of Medical Research as Emeritus Scientist. This work was supported by a Research Grant (BT/PR23486/BRB/10/1589/2017) to SK from the Department of Biotechnology, Government of India, and Research Fellowships from SCTIMST, Trivandrum, to AST and the Department of Biotechnology, Government of India, to HV. The source of funding for MW and EGL was the Intramural Research Program of the National Institute on Aging, National Institutes of Health.

## Institutional Review Board Statement

The study on rats was approved by the Institutional Animal Ethics Committees of Sree Chitra Tirunal Institute for Medical Sciences and Technology (B form No: SCT/IAEC-233/AUGUST/2017/94 and SCT/IAEC-268/FEBRUARY/2018/95).

## Informed Consent Statement

Not applicable.

## Data Availability Statement

The datasets generated during and/or analyzed during the current study are available from the corresponding author on request.

## Acknowledgments

SK is grateful to the Indian Council of Medical Research for the Emeritus Scientist position. SK, AST and HV thank Ajay Kumar R of the Rajiv Gandhi Centre for Biotechnology, Trivandrum, for providing access to the Bioruptor Facility and acknowledge the facilities provided by SCTIMST, Trivandrum.

## Conflicts of Interest

The authors declare no conflict of interest.

## Notes

### Competing Interest Statement

The authors have declared no competing interest.

